# Love at first flight: Wing interference patterns are species-specific and sexually dimorphic in blowflies (Diptera: Calliphoridae)

**DOI:** 10.1101/2020.02.18.948646

**Authors:** Nathan J. Butterworth, Thomas E. White, Phillip G. Byrne, James F. Wallman

## Abstract

Wing interference patterns (WIPs) are stable structural colours displayed on insect wings which are only visible at specific viewing geometries and against certain backgrounds. These patterns are widespread among flies and wasps, and growing evidence suggests that they may function as species- and sex-specific mating cues in a range of taxa. As such, it is expected that WIPs should differ between species and show clear sexual dimorphisms. However, the true extent to which WIPs vary between species, sexes, and individuals is currently unclear, as previous studies have only taken a qualitative approach, without considering how WIPs might be perceived by the insect. Here, we perform the first quantitative analysis of inter- and intra-specific variation in WIPs across seven Australian species of the blowfly genus *Chrysomya*. Using multispectral digital imaging and a tentative model of blowfly colour vision, we provide quantitative evidence that WIPs are species-specific, highlight that the extent of divergence is greater in males than in females, and demonstrate sexual dimorphisms in several species. These data provide evidence that WIPs have diversified substantially in blowflies and suggests that sexual selection may have played a role in this process.

## INTRODUCTION

When considering the vast suite of signals involved in animal communication, few capture the collective human interest more than those involving vision. Visual signals have been studied across an enormous variety of animal taxa, from birds (Dale et al. 2015), to frogs (Bell et al. 2017), lizards (McDiarmid et al. 2017), fish (Gerlach et al. 2014), spiders (Girard et al. 2011), and flies (White et al. 2019). Despite the breadth of this work, research continues to unravel novel modes of visual communication. Recently, there have been many discoveries of cryptic modes of visual communication – signals that are visible only to select audiences or under certain ecological settings. These inconspicuous signals are particularly prevalent among insects, most likely due to their unique and diverse visual ecologies (Lunau 2014). Examples include UV iridescent wing-spots that can only be seen from particular viewing angles (White et al. 2015), high-frequency wing-flashes that require rapid visual processing to be perceived (Eichorn et al. 2017), and colourful thin-film wing interference patterns (WIPs) that only appear at specific geometries and against certain backgrounds (Shevstova et al. 2011; Katayama et al. 2014).

WIPs are particularly widespread, and are found across all Hymenoptera, Diptera, Odonata, and some Hemiptera (Shevstova et al. 2011; Simon 2013; Brydegaard et al. 2018). They appear as brilliant patterns of colour that span the entire wing and are caused by the same process that leads to the array of colours seen in bubbles of soap. This process is referred to as two-beam thin film interference, and is caused by the interaction between light and the chitinous wing membrane. The specific geometry, hue, and intensity of insect WIPs is dependent on several variable aspects of wing morphology, including: 1) membrane thickness, since areas of differing thickness will reflect different interference colours, 2) wing corrugation, which scatters light in a coherent manner and determines the angle of interference reflection, and 3) the placement of michrotrichia, which produces spherical reflection around the base of each hair, resulting in a more ‘pebbled’ WIP appearance (Shevstova et al. 2011). Importantly, while WIPs remain stable over the lifespan of individuals (and even long after death), they exhibit limited-view iridescence, whereby the visibility of the pattern diminishes at acute geometries and against certain backgrounds (Shevstova et al. 2011).

While it is well known that many insect taxa possess exceptional vision and are capable of perceiving and discriminating colours (Hymenoptera: Peitsch et al. 1992; Diptera: Lunau 2014), the biological function of WIPs has long been overlooked. However, a growing body of research suggests that they may function as species- and sex-specific mating cues across a wide range of insects. In support of this, WIPs have been reported to be qualitatively species-specific across many Diptera (Shevstova et al. 2011), Hymenoptera (Buffington and Sandler 2011; Shevtsova and Hansson 2011), and Hemiptera (Simon 2013) – including between closely related species. There is also direct evidence that WIPs play an important role in sexual behaviour, as they have been correlated with male mating success and shown to evolve in response to sexual selection in *Drosophila* species (Katayama et al. 2014; Hawkes et al. 2019).

Despite this apparent role in reproduction, WIPs have been studied in less than 0.01% of insects – and there have been no attempts to quantitatively assess inter- and intra-specific variation. Most previous comparative studies have only approached WIP analysis from a qualitative perspective, without statistical interpretation, and without considering how WIPs are perceived by the viewer (Buffington and Sandler 2011; Shevstova et al. 2011; Shevstova and Hansson 2011; Simon 2013). Furthermore, of the few studies that have quantitatively measured WIPs, none have explicitly tested whether WIPs are species-specific or sexually dimorphic (Katayama et al. 2014; Brydegaard et al. 2018; Hawkes et al. 2019). As such, our current understanding of how WIPs vary between species, sexes, and individuals, is lacking. To address this, there is a need for studies that quantify inter- and intra-specific variation across a range of taxa, particularly in a quantitative and viewer-dependent context. Such comparative studies are necessary for informing hypotheses regarding the biological function of WIPs, while also serving as a quantitative basis for the use of WIPs in insect taxonomy.

The blowflies (Diptera: Calliphoridae) provide an ideal system to investigate the diversity and function of WIPs. Blowflies possess exceptional visual acuity and colour vision (Kirschfield 1983; van Hateren et al. 1989; Lunau 2014), and many species rely heavily on visual cues for sexual communication (Jones et al. 2014; Eichorn et al. 2017; Butterworth et al. 2019). These characteristics are especially apparent in the genus *Chrysomya*, in which many species exhibit sexually dimorphic eye morphology, in the form of holoptic eyes and ocular ‘bright zones’ in males (van Hateren et al. 1989), which are presumably involved in the recognition of light-based mating signals. Further to this, vision appears to play an important role in the sexual behaviour of two Australian species; *Ch. varipes* (Jones et al. 2014) and *Ch. flavifrons* (Butterworth et al. 2019). Here, we address this topic by quantitatively assessing the inter-and intra-specific variation of WIPs across seven species of Australasian *Chrysomya*. Considering their heavy reliance on visual signals in mate choice and recognition, and the diversity of their sexual behaviour we predict that WIPs will be highly species-specific and sexually dimorphic in this genus.

## METHODS

### Flies

Wild flies of seven species of Australian *Chrysomya* (*Ch. rufifacies, Ch. incisuralis, Ch. varipes, Ch. flavifrons, Ch. megacephala, Ch. saffranea*, and *Ch. semimetallica*) were hand netted over carrion bait between Wollongong, NSW and Brisbane, Queensland between October 2018 and March 2019. A total of 10 - 20 adults of each sex were collected, euthanised, and brought back to the lab at the University of Wollongong. Both left and right wings were removed from each fly and suspended between a glass slide and coverslip to be later photographed, for a total of 413 wings. As flies age, substantial damage and fraying occurs along the wing margin, and out of the 413 wings retrieved from wild specimens, only 231 were suitably intact for imaging and analysis.

### Photos

Wings were mounted with transparent UHU glue onto a custom rotating stage and positioned at a 45° angle which maximised WIP visibility. Photos were taken of both the left and right wing of each fly with a MZ16A stereomicroscope mounted with a Leica DFC295 digital microscope colour camera. All photos were taken at the same magnification, under standardised and uniformly diffuse lighting provided by a Leica LED5000 HDI illuminator. The Leica DFC295 produces non-linear images (in the visible spectrum), which are unsuitable for objective measurement (Hawkes et al. 2019). As such, we processed our whole-wing images using the Multispectral Image Analysis and Calibration Toolbox for ImageJ (MICA toolbox) (Troscianko et al. 2019). This produces linearized, calibrated images which allow for the measurement of relative reflectances. We calibrated our images against a 3% reflectance standard from an X-rite colour checker passport, which was placed 5 mm below the wing in the background of each photo. This resulted in a total of 231 multispectral images (visible spectrum only) of left and right wings across the seven *Chrysomya* species.

From these multispectral images, we were able to take measurements of the average values of red, green, and blue (RGB) channels (hereafter referred to as mean ‘colour’) and the standard deviation in RGB (hereafter referred to as ‘colour contrast’) across five individual wing cells (Figure 1) as well as a measurement of the entire wing. Based on these measurements, wing cells that consisted of a single colour (i.e. only red) would have a high mean colour, but low contrast, while wing cells that consisted of several colours would have high contrast (Hawkes et al. 2019). In addition to this viewer-independent analysis, we used a cone-mapping approach to convert the multispectral images into two viewer-subjective formats; the CIELab model of human colour sensation, and a receptor-based model of ‘blowfly vision’ based on the visual phenotype of *Calliphora*. Using these different models (RGB, CIELab, blowfly) we were able to assess the robustness of our results across three independent datasets. CIELab is a perceptually uniform model of human vision, whereby ‘L’ represents lightness, ‘a’ represents values on a green-red axis, and ‘b’ represents values on a blue-yellow axis. We measured the average L, a, and b pixel values (hereafter referred to as human ‘colour’) and standard deviation in L, a, and b pixel values (hereafter referred to as human ‘colour contrast’). The CIELab model allowed us to validate whether human-perceived qualitative differences in WIPs translate to quantitative differences – which will be important for their use in insect taxonomy. For the blowfly visual model, we were unable to measure UV reflectance due to the limitations of our digital microscope camera. As such, we created a simple receptor-based model of blowfly colour vision, based on the long-wavelength sensitivities of *Calliphora* (Kirschfield 1983; Hardie and Kirschfield 1983), as there are no published receptor sensitivities for *Chrysomya* species. We assumed involvement of the R8p (Rh5 opsin) and R8y (Rh6 opsin) receptors, which partly mediate colour vision (Lunau 2014), as well as the R1-6 receptors (Rh1 opsin) which contribute to both colour and luminance vision in flies (Schnaitmann et al. 2013). We estimated the mean quantum catch of Rh5, Rh6 and Rh1 (hereafter blowfly ‘colour’) as well as their standard deviation (hereafter blowfly ‘colour contrast’) across each of five individual wing cells, as well as the entire wing. This blowfly model allowed us to assess WIP variation in the context of the most ecologically relevant viewer, and the likely agent of selection on these patterns.

**Figure 1.**
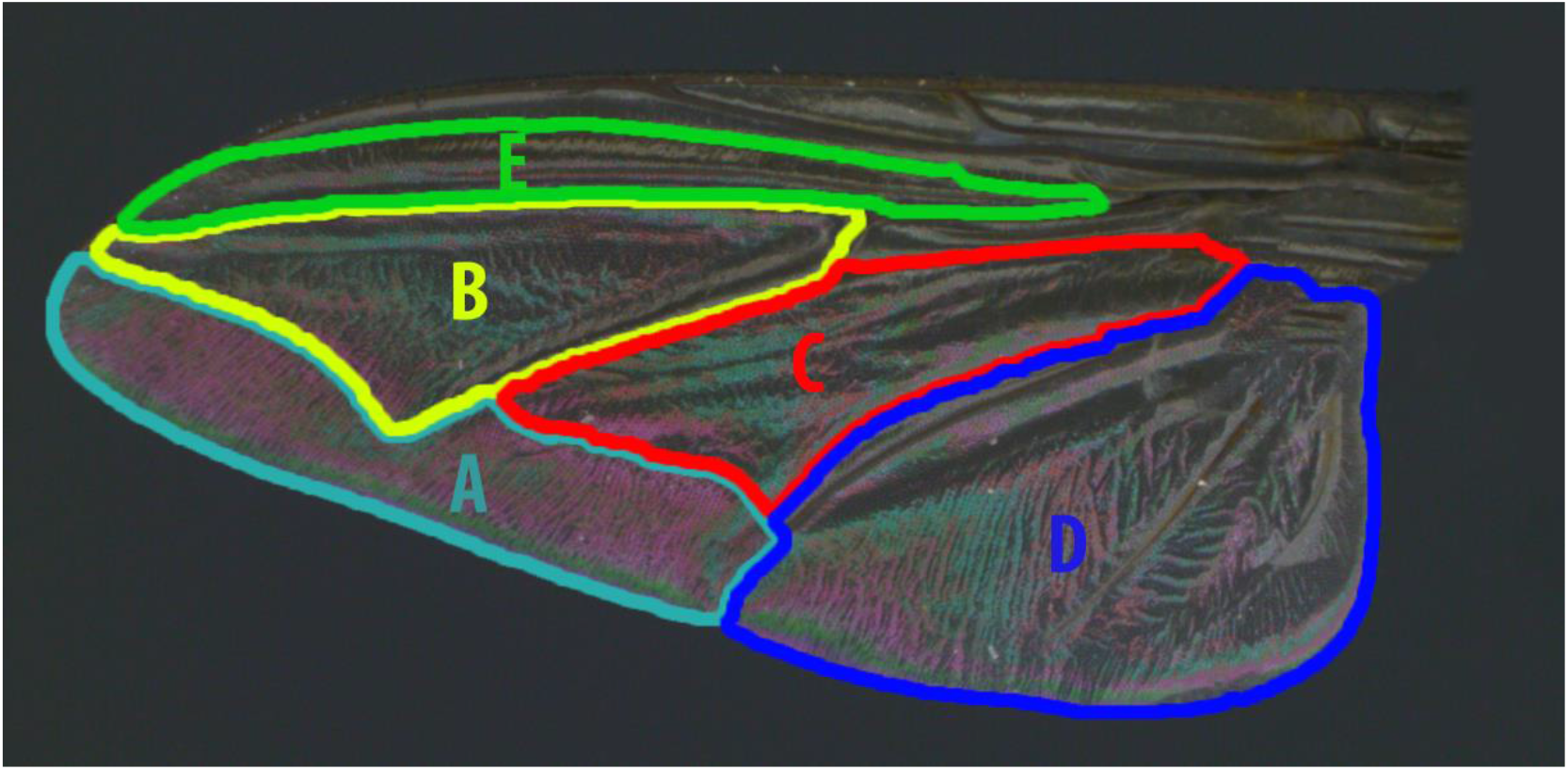
The five wing cells used for mean and standard deviation measurements of WIP colour and colour contrast across seven *Chrysomya* species. Wing cells denoted are A: 2^nd^ posterior, B: radial 4 + 5, C: discal medial, D: anterior cubital, E: radial 2 + 3. Measurements were made for RGB, CIELab, and blowfly colour space. Measurements of the whole wing were also made.

### Statistical analysis

To broadly assess the patterns of variation in the wing interference patterns of Australian *Chrysomya*, we first assessed the effects of species, sex, and wing side (left or right) on WIP variation. To do this, we first added a small constant (0.1) to each dataset (RGB, CIELab, and blowfly) to remove zeros associated with damaged wing-sections that were not measured. We then scaled each dataset using the inbuilt R scale function (R Core Team 2019) and performed a redundancy discriminant analysis (RDA) on each using the R packages ‘vegan’ (Oksanen et al. 2019) and ‘RVAideMemoire’ (Hervé 2019). To validate the effect of species, sex, and wing on WIP variation, the total percentage of constrained variance explained by the three factors was estimated by a canonical *R*^2^ called the ‘bimultivariate redundancy statistic’ (Miller and Farr 1971; Peres-Neto et al. 2006; Hervé et al. 2018). For the RGB, CIELab, and blowfly datasets species, sex, wing, and their interactions explained 46% (RGB), 38% (CIELab), and 51% (Blowfly) of the total variation in WIP colour and 62% (RGB), 58% (CIELab), and 53% (Blowfly) of the total variation in WIP colour contrast. To test whether these constrained variances constituted a significant proportion of the variation in each dataset, permutation *F*-tests based on the canonical *R*^2^ were performed (Legendre and Legendre 2012; Hervé et al. 2018). The tests were all declared significant (PERMANOVA; P < 0.001), which implies that the chosen factors (species, sex, and wing) explained a significant proportion of the total variation in colour and contrast in each of the three datasets. As such, to test for the individual effects of each factor, a second permutation *F*-test was performed for species, sex, wing and the species × sex × wing interaction.

To assess the differences between species while accounting for sex-specific variance, we separated the CIELab and blowfly datasets into male and female datasets and performed two further RDAs. For these analyses, we used only measurements from the left wings, as preliminary inspections showed asymmetries between left and right wings within species (Figures S1 & S2). For the female datasets, species explained 34% (CIELab) and 51% (Blowfly) of the total variation in WIP colour and 54% (CIELab) and 59% (Blowfly) of the total variation in WIP colour contrast. For the male datasets, species explained 36% (CIELab) and 45% (Blowfly) of the total variation in WIP colour and 58% (CIELab) and 47% (Blowfly) of the total variation in WIP colour contrast. To test whether these variances constituted a significant proportion of the data, permutation *F*-tests based on the canonical *R*^2^ were performed. The tests were all declared significant (PERMANOVA; P < 0.001), which implies that differences in colour and colour contrast between species explained a substantial portion of the total variation of each dataset. As such, a pairwise comparison using the function ‘pairwise.factorfit’ from ‘RVAideMemoire’ was used to specifically assess which species differed significantly from each other within the male and female datasets. Lastly, to assess intra-specific variation (i.e. whether WIPs were sexually dimorphic), datasets were separated into species, resulting in seven individual CIELab datasets and seven individual blowfly datasets. To consider the effect of sex, each dataset was scaled with the inbuilt R function, and principal component analysis (PCA) was conducted. Univariate analysis of variance (ANOVA) was then performed on the extracted PCs from each dataset to test for significant differences in PCs (representing colour or contrast) between male and female wings. All PCA and ANOVA analyses were performed using the R base package (R Core Team 2019), the ‘Factoextra’ package (Kassambra and Mundt 2017), and the ‘ggFortify’ package (Tang et al. 2016).

## RESULTS

Initial observations indicated that there was substantial inter-specific variation in WIPs, with clear differences between species. *Ch. rufifacies* and *Ch. incisuralis*, for example, showed vastly different WIPs compared to *Ch. flavifrons* and *Ch. varipes* (Figure 2). There were also noticeable intra-specific differences between male and female WIPs in both colour and colour contrast, particularly in *Ch. flavifrons* (Figure 2). Further to this, preliminary examination revealed asymmetries between left and right WIPs within individuals (Figures S1 & S2).

**Figure 2.**
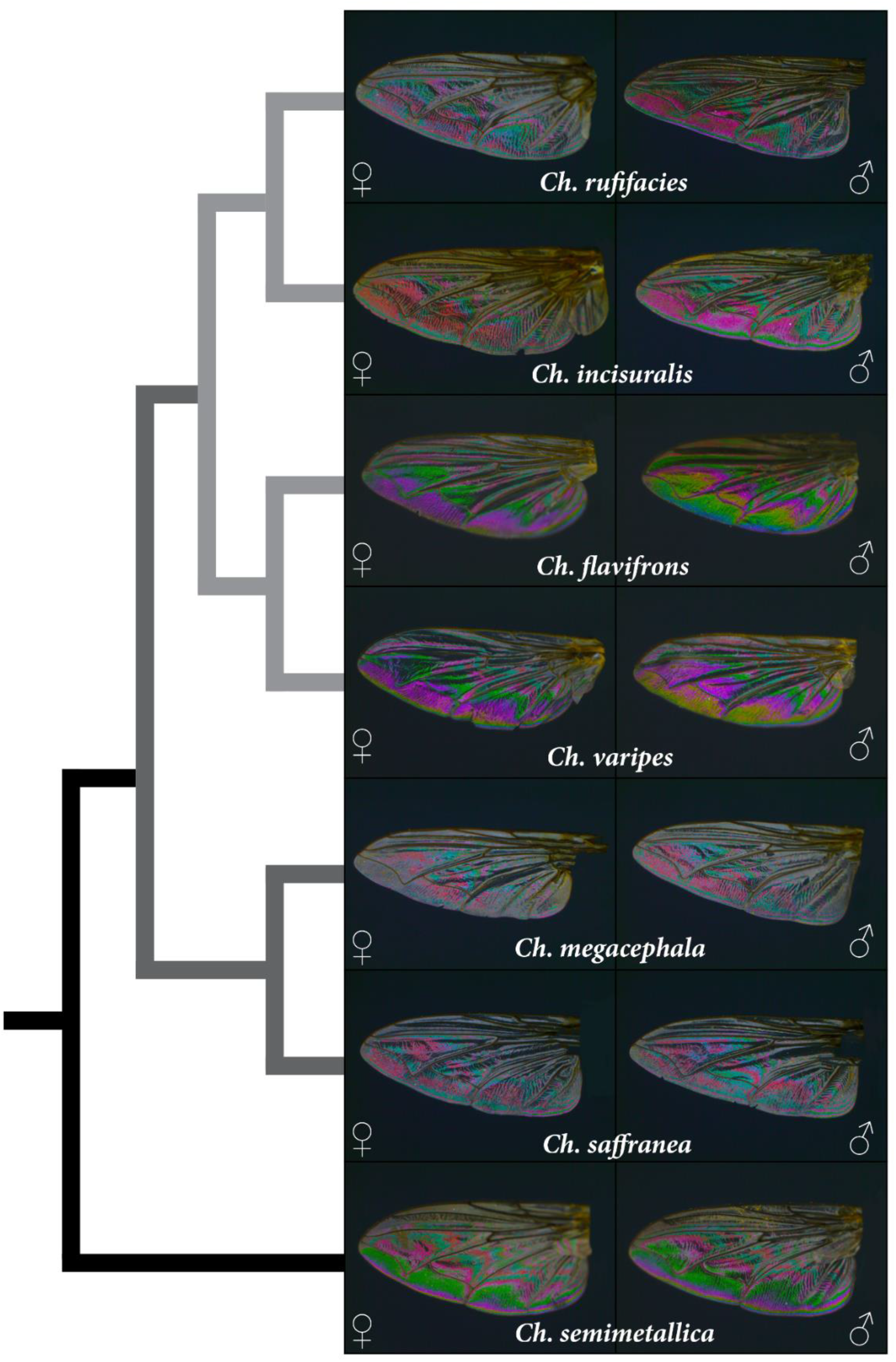
WIP variation among seven species of Australian *Chrysomya* (Diptera: Calliphoridae). Images captured with an MZ16A stereomicroscope mounted with a Leica DFC295 digital microscope colour camera. All photos were taken at the same magnification, under standardised and uniformly diffuse lighting provided by a Leica LED5000 HDI illuminator. To improve figure clarity, the contrast and saturation of each WIP were raised by 40% in Adobe Lightroom 2019. The final figure was edited with Adobe InDesign 2019. The reduced phylogeny of the seven Australian species is based on Singh et al. 2011. Clade I represented by light grey branches, Clade II by dark grey branches, and Clade III by black branches.

To assess these patterns of variation, while accounting for species, sex, and wing, RDA was performed. The RDA revealed that the combined effect of species, sex, and wing explained a significant proportion of overall variation in colour and contrast across RGB, CIELab and blowfly datasets. Of the constrained variance (the variance explained by all three factors), discriminant components 1-5 collectively accounted for 95.17% (RGB), 91.89% (CIELab), 98.10% (blowfly) of the variation in colour, and 98.04% (RGB), 97.36% (CIELab), 97.58% (blowfly) of the variation in contrast. Permutation F-tests suggested that species (PERMANOVA; P < 0.001), sex (PERMANOVA; P P < 0.001), and the species × sex interaction (PERMANOVA; P P < 0.001) each individually explained a significant proportion of colour and colour contrast variation across all three models (RGB, CIELab and Blowfly) (Table S1). While wing also explained a significant proportion of colour variation in the RGB and CIELab datasets (PERMANOVA; P P < 0.05), this was not significant when considered as an interaction with species, sex, or species × sex (Table S1). However, considering that there were asymmetries between mean values of left and right wings within species (though not statistically significant) (Figures S1 & S2) we opted to perform all subsequent analyses with left wings only.

### Inter-specific comparisons

To assess how WIPs varied between species, we had to account for the sexual variation in WIP colour and contrast. To do so, a second RDA was performed on individual male and female datasets (for CIELab and blowfly visual space). The RDA revealed substantial inter-specific variation in WIPs in both the blowfly (Figure 3) and CIELab datasets (Figure S3), whereby species explained a significant proportion of the variation in male WIP colour (CIELab: 35.74%; Blowfly: 45.24%), male WIP contrast (CIELab: 57.35%; Blowfly: 46.74%), female WIP colour (CIELab: 34.27%; Blowfly: 51.30%) and female WIP contrast (CIELab: 53.94%; Blowfly: 58.67%). Pairwise comparisons on the blowfly dataset (Table 1) showed that for females, variation in WIP colour did not separate any species from their closest relatives (Pairwise comparison: P > 0.05). However, female variation in WIP contrast clearly separated *Ch. varipes* from its sister species *Ch. flavifrons* (Pairwise comparison: P < 0.05). In males, variation in WIP colour separated all species from their closest relatives (Pairwise comparisons: P < 0.05), with the exception of *Ch. megacephala* and *Ch. saffranea* (Pairwise comparisons: P > 0.05). Similarly, male variation in WIP contrast separated all species from their closest relatives (Pairwise comparisons: P < 0.05). Pairwise comparisons of the CIELab data showed similar results, whereby variation in both WIP colour and WIP contrast significantly separated all closely related species (Pairwise comparisons: P < 0.05) (Table S2).

**Table 1.** Pairwise comparisons between species, based on redundancy discriminant analysis of WIP colour (as represented by average measurements of Rh5, Rh6, and Rh1 values) and WIP colour contrast (as represented by standard deviations in Rh5, Rh6, and Rh1 values). All measurements were made in ‘blowfly visual space’ using the receptor sensitivities of *Calliphora* in the Multispectral Image Analysis and Calibration Toolbox for ImageJ (MICA toolbox) (Troscianko et al. 2019). Bold values indicate significant differences. F = Female, M = Male.

| **F_Colour** | Fla | Inc | Meg | Ruf | Saf | Sem |
| --- | --- | --- | --- | --- | --- | --- |
| Inc | 0.0567 | - | - | - | - | - |
| Meg | 0.88 | **0.0042** | - | - | - | - |
| Ruf | 0.6363 | 0.3399 | 0.6363 | - | - | - |
| Saf | **0.042** | 0.3399 | 0.2181 | 0.2289 | - | - |
| Sem | **0.0042** | **0.042** | **0.0042** | **0.007** | **0.0042** | - |
| Var | 0.6363 | **0.009** | 0.133 | 0.5052 | 0.0745 | **0.0042** |
| **F_Contrast** | Fla | Inc | Meg | Ruf | Saf | Sem |
| Inc | 0.0952 | - | - | - | - | - |
| Meg | 0.1598 | **0.0952** | - | - | - | - |
| Ruf | **0.0382** | 0.376 | **0.0385** | - | - | - |
| Saf | **0.021** | **0.1221** | 0.1598 | 0.2719 | - | - |
| Sem | **0.0052** | 0.3392 | 0.0052 | 0.0052 | 0.006 | - |
| Var | **0.042** | **0.006** | 0.0105 | 0.006 | 0.021 | **0.0052** |
| **M_Colour** | Fla | Inc | Meg | Ruf | Saf | Sem |
| Inc | 0.38 | - | - | - | - | - |
| Meg | **0.0026** | 0.1802 | - | - | - | - |
| Ruf | **0.0026** | **0.0378** | **0.0117** | - | - | - |
| Saf | **0.0026** | **0.0134** | 0.1155 | **0.0126** | - | - |
| Sem | **0.0026** | **0.0315** | **0.0026** | 0.0692 | **0.0026** | - |
| Var | **0.0026** | **0.0158** | 0.064 | **0.0275** | 0.2604 | **0.0026** |
| **M_Contrast** | Fla | Inc | Meg | Ruf | Saf | Sem |
| Inc | **0.0338** | - | - | - | - | - |
| Meg | **0.0115** | 0.189 | - | - | - | - |
| Ruf | **0.0052** | **0.0225** | **0.0093** | - | - | - |
| Saf | **0.003** | **0.0238** | **0.0289** | **0.0105** | - | - |
| Sem | **0.003** | 0.0696 | **0.003** | **0.003** | **0.003** | - |
| Var | **0.003** | **0.014** | 0.1722 | **0.0145** | **0.0338** | **0.003** |

**Figure 3.**
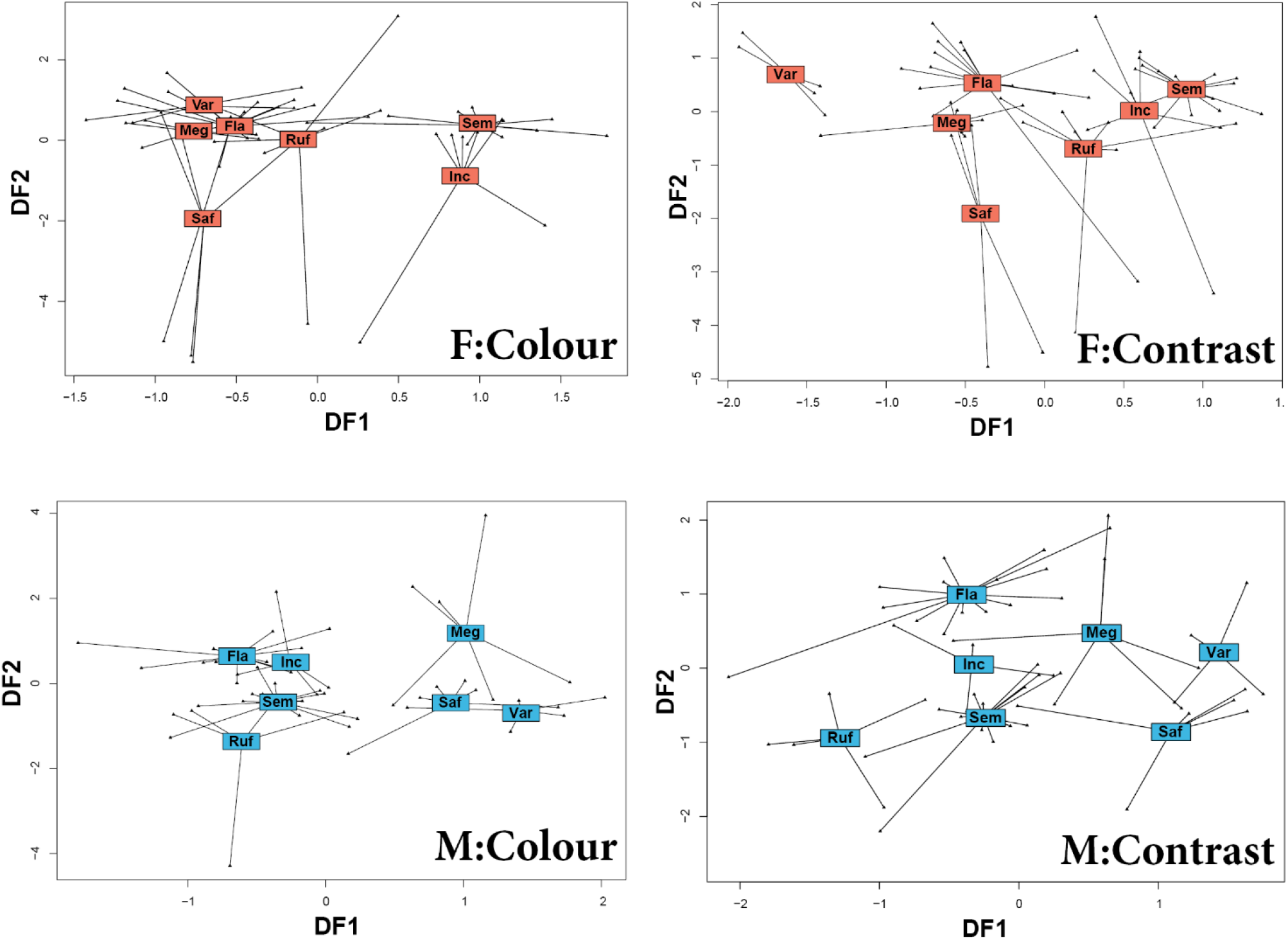
Quantitative differences in the wing interference patterns (WIPs) of male (M) and female (F) Australian *Chrysomya* represented by discriminant factors 1 (DF1) and 2 (DF2). Results are from a redundancy discriminant analysis of WIP colour (as represented by average measurements of Rh5, Rh6, and Rh1 values) and WIP colour contrast (as represented by standard deviations in Rh5, Rh6, and Rh1 values). All measurements were made in ‘blowfly visual space’ using the receptor sensitivities of *Calliphora* in the Multispectral Image Analysis and Calibration Toolbox for ImageJ (MICA toolbox) (Troscianko et al. 2019).

### Intra-specific comparisons

To investigate and visualize sex-specific differences within each of the seven species, we separated the CIELab and blowfly datasets by species. On each of these datasets PCA and univariate ANOVA were performed, revealing quantitative sexual dimorphisms in the blowfly data in WIP colour (Figure 4) and colour contrast (Figure 5) for several *Chrysomya* species. Similar patterns were observed in the CIELab datasets (Figure S4 & Figure S5). Of these sex-specific differences, the first five PCs explained a substantial proportion (>80%) of the overall variation in WIP colour and contrast in both the CIELab and blowfly datasets (Tables S3-a, S4-a, S5-a, S6-a). As such, ANOVA was performed on the first five PCs extracted from these datasets for each species. For the blowfly data, this revealed significant differences between male and female WIP colour in *Ch. rufifacies, Ch. flavifrons*, *Ch. megacephala* and *Ch. semimetallica* (Table S3-a). Further, WIP contrast also showed sex-specific differences in *Ch. rufifacies*, *Ch. flavifrons*, and *Ch. varipes* (Table S4-a). Similarly, the first five PCs extracted from the CIELab dataset showed sex-specific differences in WIP colour and contrast for all the above species, as well as for *Ch. saffranea* (Tables S5-a & S6-a). To determine which variables (i.e. which aspects of colour and which wing cells) contributed to each principal component, we used the ‘fviz_contrib’ function from ‘factoextra’. To see which variables characterise the sexual differences in WIP colour and contrast for each of the seven *Chrysomya* species, see Tables S3-b, S4-b, S5-b and S6-b.

**Figure 4.**
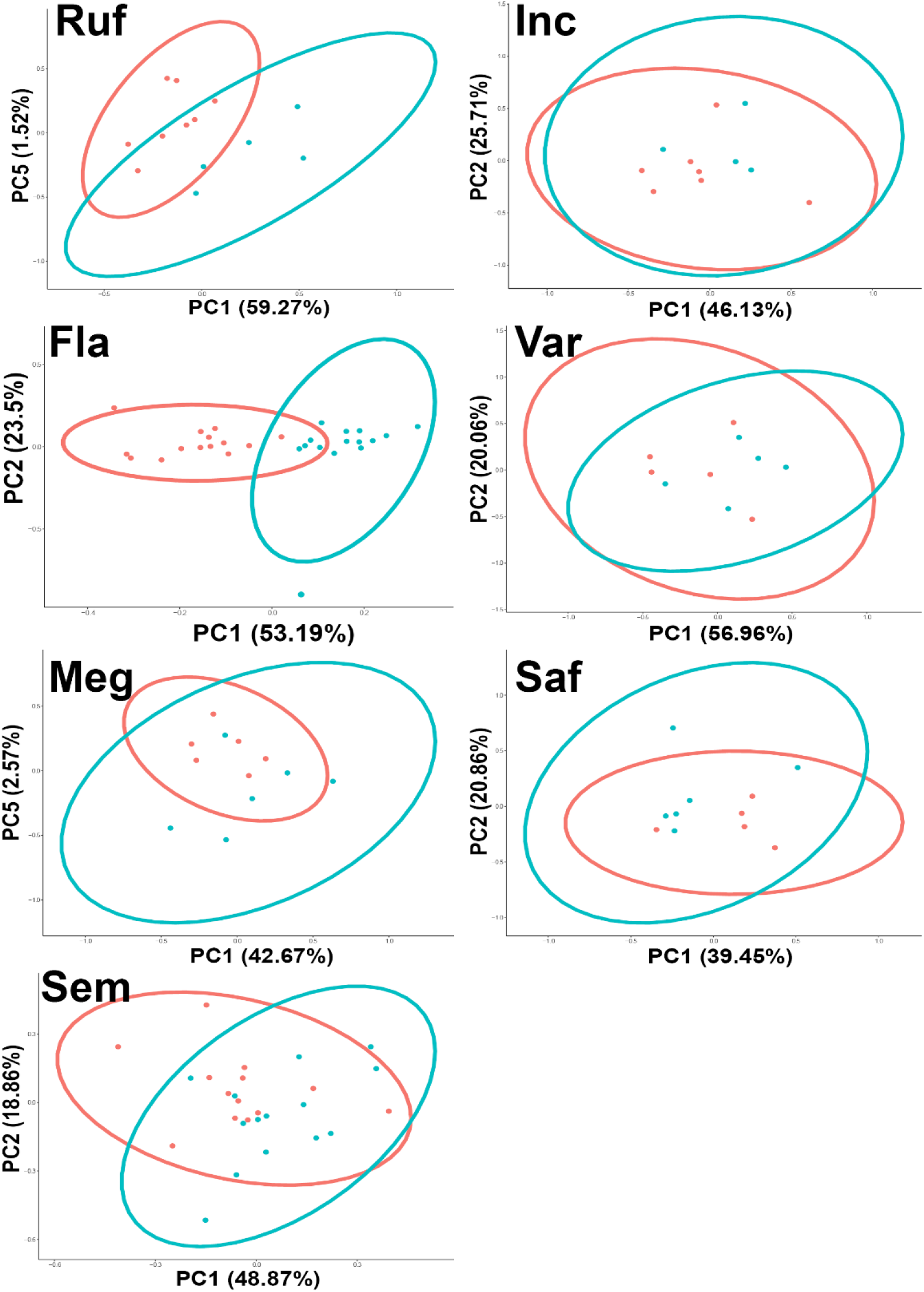
PCA plots of sex-specific differences in the ‘blowfly’ average colour of WIPs (mean Rh1, Rh5 and Rh6 values). The blue dots and ellipses represent males, while red dots and ellipses represent females. All measurements were made in ‘blowfly visual space’ using the receptor sensitivities of *Calliphora* in the Multispectral Image Analysis and Calibration Toolbox for ImageJ (MICA toolbox) (Troscianko et al. 2019).

**Figure 5.**
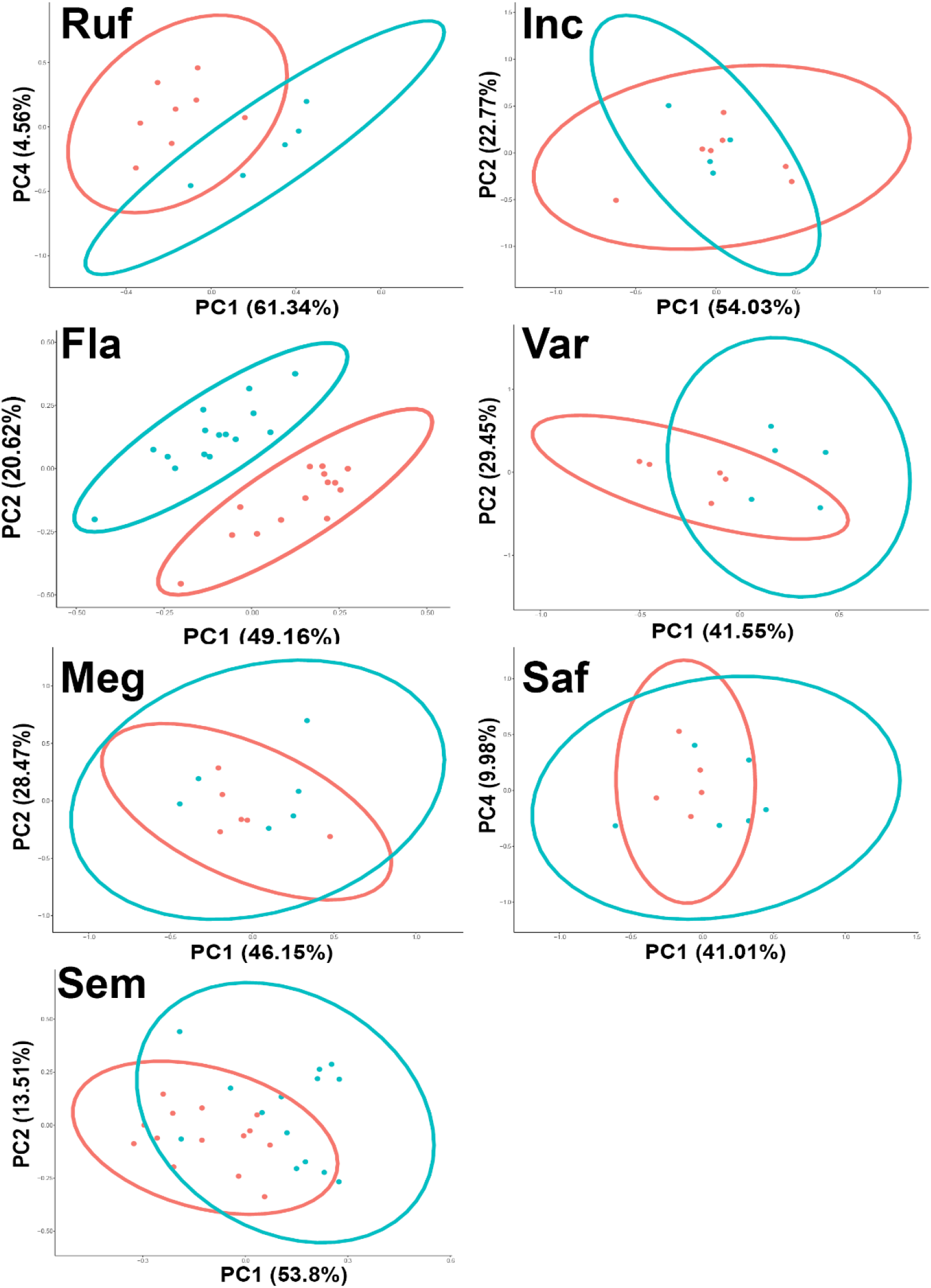
PCA plots of sex-specific differences in the ‘blowfly’ colour contrast of WIPs (standard deviation in Rh1, Rh5 and Rh6 values). The blue dots and ellipses represent males, while red dots and ellipses represent females. All measurements were made in ‘blowfly visual space’ using the receptor sensitivities of *Calliphora* in the Multispectral Image Analysis and Calibration Toolbox for ImageJ (MICA toolbox) (Troscianko et al. 2019).

## DISCUSSION

Wing interference patterns are widespread among insects, and accumulating evidence suggests that they may function as species- and sex-specific mating cues. Despite this, past inter- and intra-specific comparisons have been limited to qualitative assessments. Here, we provide quantitative evidence that WIPs are species-specific in the blowfly genus *Chrysomya*. We also show that the extent of divergence is greater in males than in females, and highlight significant sexual dimorphisms in several species. Our findings support the notion that WIPs may play an important role in blowfly mating behaviour by functioning as species- and sex-specific mating cues.

### Species differences

Since the RGB, CIELab, and blowfly analyses all produced qualitatively similar results, the subsequent discussion will focus primarily on the results of the blowfly-based analyses, as these data represent the most ecologically relevant receiver. Our results highlight substantial diversification in WIPs in *Chrysomya*, with significant differences between several species, particularly between close relatives. Notably, the patterns of inter-specific variation differed between males and females; female differences in WIP colour (that is the average colour as measured in our blowfly model) did not separate close relatives, whereas female differences in WIP contrast (that is the number of contrasting colours as measured in our blowfly model) clearly separated female *Ch. varipes* from *Ch. flavifrons*. In males, divergence between species was greater, whereby the WIPs of most closely related species diverged substantially. For example, WIP colour separated *Ch. incisuralis* from *Ch. rufifacies*, and *Ch. varipes* from *Ch. flavifrons*, while WIP contrast separated *Ch. saffranea* from *Ch. megacephala*. These differences were even more pronounced in the CIELab data (Table S2), where almost every species separated based on WIP colour and WIP contrast. However, *Ch. megacephala* and *Ch. saffranea* overlapped substantially in both the blowfly and CIELab datasets, indicating limited divergence in WIPs between these two very closely related species. Further to this, there was substantial overlap in both blowfly and CIELab measurements between the *Ch. megacephala*/*Ch. saffranea* species group and the distantly related *Ch. incisuralis*/*Ch. rufifacies* species group, which suggests convergent evolution in WIP patterns in these two groups.

Our data also suggest that selection for WIP divergence differs between males and females. For example, *Ch. incisuralis* and *Ch. rufifacies* males differ based on WIP colour and WIP contrast, while females do not differ in either measurement. Likewise, males of *Ch. saffranea* and *Ch. megacephala* differ in WIP colour contrast, but females do not differ in either measurement. Moreover, males of *Ch. varipes* and *Ch. flavifrons* differ in WIP colour and WIP contrast, while females only differ in WIP contrast. If blowfly WIPs are in fact used as mating cues, these results might suggest that WIP divergence is primarily driven by selection on male wings. This is supported by findings from previous work in *Drosophila* species, where male WIPs, but not female WIPs, have been shown to experience sexual selection (Hawkes et al. 2019). Importantly, when comparing between males of different species (except *Ch. saffranea* and *Ch. megacephala*) it was both the mean colour and colour contrast of WIPs that varied – suggesting that both aspects of the pattern may be relevant in the context of signalling. This is supported by findings in *Drosophila simulans* where there was evidence for sexual selection on average wing colour, colour contrast, as well as luminance, across the whole wing (Hawkes et al. 2019). As such, both the average colour of the WIP, and the number of contrasting colours within, are likely to be important aspects of fly WIPs, and future studies should consider both traits when making comparisons.

It is also plausible that the species-specific differences in WIPs we report are unrelated to sexual selection but are instead a side effect of differences in body size and wing morphology between species. This is because body size and wing membrane thickness tend to scale allometrically (Wootton 1992) which has a direct effect on the colours reflected in WIPs. Specifically, the sequence of WIP colours corresponds to the Newton series reflected from a thin film of oil on water (Shevstova et al. 2011; Katayama et al. 2014). The first three Newton orders (0 to 550 nm wing membrane thickness) are the brightest and display a near complete scale of spectral colours, except for pure red. This explains why the smaller species, *Ch. varipes*, *Ch. flavifrons*, and *Ch. semimetallica* (~3-6 mm body length), with thinner wing membranes show brighter WIPs composed of blues, greens, yellows, and purples (Figure 2). Conversely, larger species with thicker wing membranes (≥550 nm wing membrane thickness) appear to display duller WIPs (Buffington and Sandler 2011) composed of non-spectral (to the human eye) magentas and greens that gradually fade into uniform pale grey. This is apparent in the larger *Chrysomya* species (*Ch. incisuralis*, *Ch. rufifacies*, *Ch.megacephala* and *Ch. saffranea*; all ~8-12 mm body length) and explains why the WIPs of these species overlap substantially. Therefore, the substantial differences between the species pairs *Ch. varipes/Ch. flavifrons* and *Ch. incisuralis/Ch rufificacies* can be primarily attributed to gross differences in body size and wing membrane thickness.

While larger blowfly species tended to display duller WIPs, the differences in colour patterns are still statistically distinct in our model of blowfly colour vision, separating *Ch. rufifacies* and *Ch. incisuralis* across several measurements. Therefore, it is plausible that even the duller WIPs of larger blowflies may still act as species- and sex-specific cues. Gross differences in body size cannot, however, explain the observed divergence in WIPs between species with similar body and wing sizes. For example, male WIPs of *Ch. incisuralis* and *Ch. rufifacies* clearly diverge, but body and wing size are almost identical in both species. Likewise, in *Ch. varipes* and *Ch. flavifrons*, stark differences in WIPs are apparent between females of both species, even though they exhibit similar wing structure (Aldrich 1925). Therefore, the differences in WIPs between these closely related species must be due to more fine-scale differences in wing membrane thickness, perhaps restricted to specific parts of the wing. While these fine-scale, species-specific differences in wing structure may result from sexual selection on WIPs as species- and sex-specific signals, it is also likely that they are the result of differing ecological selection on wing morphology for flight performance (Taylor and Merriam 1995; DeVries et al. 2010).

### Sex differences

If sexual selection has acted on the WIPs of male *Chrysomya*, then we might expect to see evidence of sexual dimorphism, either in WIP colour or colour contrast, across multiple species. Correspondingly, sexual dimorphism in PCs were apparent for five of the seven species. *Ch. rufifacies, Ch. flavifrons*, *Ch. megacephala*, and *Ch. semimetallica* all showed sex-specific differences in the average colour and contrast of WIPs. Whereas *Ch. varipes* only showed sex-specific differences in WIP colour contrast. Importantly, while the whole wing contributed to the sexual variation of some species, in most species it was specific wing cells that contributed most of the sex-specific variation (Table S3-b). This suggests that certain sections of the wing may be under stronger selection than others, and highlights that taking measurements across the whole wing can in fact cloud patterns of inter- and intra-specific variation. The use of highly localised colour patterns as signals has been demonstrated in many other animal taxa (Breuker and Brakefield 2002; Fleishman et al. 2017) and may partly explain why no sexual dimorphism was apparent across the whole wing measurements of *Drosophila simulans* (Hawkes et al. 2019).

The greatest degree of sexual dimorphism observed in the present study was in *Ch. flavifrons* – a species where visual cues are known to play a key role in mating behaviour during male courtship displays (Butterworth et al. 2019). This was predominantly driven by differences in the average colour of wing cell E, and the colour contrast of wing cells B and C. The sex-specific differences in the average colour of wing cell E are likely due to the fumosity (light brown pigmentation) extending from the wing margin of males, which is not present in females. Pigmentation is known to substantially affect interference colouration, likely constituting an important component of WIP displays in numerous flies and wasps (Shevstova et al. 2011) and has likely evolved as a component of the male courtship display in *Ch. flavifrons* (Butterworth et al. 2019). Nevertheless, sexual dimorphism was also observed in wing cells B and C of *Ch. flavifrons*, areas where no wing pigmentation is apparent. Likewise, sexual dimorphism was apparent in *Ch. rufifacies* and *Ch. semimetallica*, two species where neither male nor female wings exhibit pigmentation. These sex-specific differences must therefore be the result of minor differences in wing membrane thickness and corrugation, both of which may be the result of selection for sex-specific WIPs.

While sexual dimorphism is often the result of sexual selection, there are also numerous examples of sexual dimorphism being driven primarily by ecological selection (Slatkin 1984; Taylor et al. 2019). For example, sexually dimorphic wing morphology resulting from sex-specific selection on flight performance has been demonstrated in *Morpho* butterflies (DeVries et al. 2010). Similarly, flight performance is known to differ between male and female blowflies, as males are adapted to chase females mid-flight (Trischler et al. 2010). The necessity for males to track females, and rapidly adjust their trajectory during flight may therefore impose selective pressure on male wing morphology, which might not be experienced by females - hence leading to sexually dimorphic membrane thicknesses and WIPs, which are unrelated to signalling. However, it seems unlikely that selection for flight performance would only result in minor changes to wing membrane thickness between the sexes, without more substantial differences in wing shape and size as is the case in *Morpho* butterflies (DeVries et al. 2010). Overall, we suggest that these differences may be primarily driven by sexual selection, particularly in *Ch. varipes* and *Ch. flavifrons;* two species where males perform complex courtship displays (Jones et al. 2014; Butterworth et al. 2019). These displays mirror those seen in *Drosophila* species, where WIPs almost certainly constitute an important component of the display (Katayama et al. 2014; Hawkes et al. 2019).

### Conclusions

In their comprehensive review of fly vision, Lunau et al. (2014) stated “Interestingly, only a few flies exhibit a dimorphism of coloured courtship signals, indicating that courtship and mating are based on cues other than colour”. Here, we provide quantitative evidence that WIPs are sexually dimorphic and differ substantially between closely related blowflies. This, in line with the recent findings that WIPs are under sexual selection in *Drosophila*, suggests that colour may play a greater role in fly mating behaviour than previously thought, and further substantiates WIPs as a promising avenue for research into colour-based mating signals in flies.

However, the study of insect WIPs is still in its infancy, and while our results show substantial species- and sex-specific differences in the WIPs of Australian *Chrysomya* – it is unclear whether these patterns extend to other taxa, and whether they are driven by ecological selection on wing morphology or sexual selection on WIP appearance. Our findings should also be tempered by the fact that we used a tentative model of blowfly colour vision, and were unable to consider UV reflectance, which may also form an important part of WIP displays – although, evidence in *Drosophila simulans* suggests that UV may play only a minor role (Hawkes et al. 2019). Furthermore, although we have demonstrated sexual dimorphisms in several parts of the wing, we used standardised and diffused lighting and a uniform background – so exactly how these differences appear to blowflies in a natural setting remains unknown. In fact, there have been no studies of WIPs under ecologically relevant settings for any species, so there is still much to learn about which aspects of the WIP are displayed and perceptible to flies under field conditions. Lastly, there is a compelling need for more studies that combine multispectral imaging, a viewer-dependent model of analysis, and behavioural assays as per Hawkes et al. (2019). We suggest that *Ch. flavifrons* will be a good candidate for such studies in blowflies.

## ACKNOWLEDGEMENTS

This project was supported by The Holsworth Wildlife Research Endowment & The Ecological Society of Australia. We thank Tracey Gibson and Natasha Mansfield for their contributions. NJB would also like to acknowledge Finlay Davidson and Sean Pryor for their support.

## REFERENCES

Aldrich JM (1925) New Diptera or two-winged flies in the United States National Museum.Proceedings of the United States National Museum 66:1–36 https://doi.org/10.5479/si.00963801.66-2555.1

Bell RC, Webster GN, Whiting M.J. (2017) Breeding biology and the evolution of dynamic sexual dichromatism in frogs. Journal of Evolutionary Biology 30:2104–2115 doi:10.1111/jeb.13170

Breuker CJ, Brakefield PM (2002) Female choice depends on size but not symmetry of dorsal eyespots in the butterfly *Bicyclus anynana*. Proceedings of the Royal Society B: Biological Sciences 269:1233–1239 doi:10.1098/rspb.2002.2005

Brydegaard M, Jansson S, Schulz M, Runemark A (2018) Can the narrow red bands of dragonflies be used to perceive wing interference patterns? Ecology and Evolution 8:5369–5384 doi:10.1002/ece3.4054

Buffington ML, Sandler RJ (2012) The occurrence and phylogenetic implications of wing interference patterns in Cynipoidea (Insecta: Hymenoptera). Invertebrate Systematics 25:586–597

Butterworth NJ, Byrne PG, Wallman JF (2019) The blow fly waltz: Field and laboratory observations of novel and complex dipteran courtship behavior. Journal of Insect Behavior 32:109–119 doi:10.1007/s10905-019-09720-1

Dale J, Dey CJ, Delhey K, Kempenaers B, Valcu M (2015) The effects of life history and sexual selection on male and female plumage colouration. Nature 527:367–370 doi:10.1038/nature15509

DeVries PJ, Penz CM, Hill RI (2010) Vertical distribution, flight behaviour and evolution of wing morphology in *Morpho* butterflies. Journal of Animal Ecology 79:1077–1085 doi:10.1111/j.1365-2656.2010.01710.x

Eichorn C, Hrabar M, Van Ryn EC, Brodie BS, Blake AJ, Gries G (2017) How flies are flirting on the fly. BMC Biology 15:1–9 doi:10.1186/s12915-016-0342-6

Fleishman LJ, Yeo AI, Perez CW (2017) Visual acuity and signal color pattern in an *Anolis* lizard. The Journal of Experimental Biology 220:2154 doi:10.1242/jeb.150458

Gerlach T, Sprenger D, Michiels NK (2014) Fairy wrasses perceive and respond to their deep red fluorescent coloration. Proceedings of the Royal Society B: Biological Sciences 281:20140787 doi:10.1098/rspb.2014.0787

Girard MB, Kasumovic MM, Elias DO (2011) Multi-modal courtship in the peacock spider, *Maratus volans* (O.P.-Cambridge, 1874). PLoS One 6:e25390 doi:10.1371/journal.pone.0025390

Hardie RC, Kirschfeld K (1983) Ultraviolet sensitivity of fly photoreceptors R7 and R8: Evidence for a sensitising function. Biophysics of Structure and Mechanism 9:171–180 doi:10.1007/BF00537814

Hawkes MF, Duffy E, Joag R, Skeats A, Radwan J, Wedell N, Sharma MD, Hosken DJ, Troscianko J (2019) Sexual selection drives the evolution of male wing interference patterns Proceedings of the Royal Society B: Biological Sciences 286:20182850 doi:10.1098/rspb.2018.2850

Hervé MR, Nicolè F, Lê Cao K-A (2018) Multivariate analysis of multiple datasets: A practical guide for chemical ecology. Journal of Chemical Ecology 44:215–234 doi:10.1007/s10886-018-0932-6

Hervé MR (2020) RVAideMemoire: Testing and plotting procedures for biostatistics. R package version 0.9–74. https://cran.r-project.org/package=RVAideMemoire

Jones SD, Byrne PG, Wallman JF (2014) Mating success is predicted by the interplay between multiple male and female traits in the small hairy maggot blowfly. Animal Behaviour 97:193–200 doi:10.1016/j.anbehav.2014.09.022

Kassambra A, Mundt F (2017) factoextra: Extract and visualize the results of multivariate data analyses. R Package Version 1.0.4. https://CRAN.R-project.org/package=factoextra

Katayama N, Abbott JK, Kjærandsen J, Takahashi Y, Svensson EI (2014) Sexual selection on wing interference patterns in *Drosophila melanogaster*. Proceedings of the National Academy of Sciences of the United States of America 111:15144–15148 doi:10.1073/pnas.1407595111

Kirschfeld K, Feiler R, Hardie R, Vogt K, Franceschini N (1983) The sensitizing pigment in fly photoreceptors. Biophysics of Structure and Mechanism 10:81–92 doi:10.1007/BF00535544

Koshio C, Muraji M, Tatsuta H, Kudo S-I (2007) Sexual selection in a moth: effect of symmetry on male mating success in the wild. Behavioral Ecology 18:571–578 doi:10.1093/beheco/arm017

Legendre P, Legendre, LF (2012) Numerical Ecology Elsevier, Amsterdam, The Netherlands

Lunau K (2014) Visual ecology of flies with particular reference to colour vision and colour preferences. Journal of Comparative Physiology A 200:497–512 doi:10.1007/s00359-014-0895-1

McDiarmid CS, Friesen CR, Ballen C, Olsson M (2017) Sexual coloration and sperm performance in the Australian painted dragon lizard, *Ctenophorus pictus*. Journal of Evolutionary Biology 30:1303–1312 doi:10.1111/jeb.13092

McLachlan AJ (2010) Fluctuating asymmetry in flies, what does it mean? Symmetry 2:1099–1107 doi:10.3390/sym2021099

Miller JK, Farr SD (1971) Bimultivariate redundancy: A comprehensive measure of interbattery relationship. Multivariate Behavioral Research 6:313–324 doi:10.1207/s15327906mbr0603_4

Møller AP, Pomiankowski A (1993) Fluctuating asymmetry and sexual selection. Genetica 89:267 doi:10.1007/BF02424520

Oksanen J, Blanchet G, Friendly M, Kindt R, Legendre P, McGlinn D, Minchin PR, O’Hara RB, Simpson GL, Solymos P, Stevens MHH, Szoecs E and Wagner H (2019) vegan: Community ecology package. R package version 2.5-5. https://CRAN.R-project.org/package=vegan

Peitsch D, Fietz A, Hertel H, de Souza J, Ventura DF, Menzel R (1992) The spectral input systems of hymenopteran insects and their receptor-based colour vision. Journal of Comparative Physiology A 170:23–40 doi:10.1007/BF00190398

Peres-Neto PR, Legendre P, Dray S, Borcard D (2006) Variation partitioning of species data matrices: Estimating and comparison of fractions. Ecology 87:2614–2625 doi:10.1890/0012-9658(2006)87[2614:VPOSDM]2.0.CO;2

R Core Team (2019). R: A language and environment for statistical computing. R Foundation for Statistical Computing, Vienna, Austria. URL https://www.R-project.org/.

Schnaitmann C, Garbers C, Wachtler T, Tanimoto H (2013) Color discrimination with broadband photoreceptors. Current Biology 23:3275–2382 doi:10.1016/j.cub.2013.10.037

Shevtsova E, Hansson C (2011) Species recognition through wing interference patterns (WIPs) in *Achrysocharoides* Girault (Hymenoptera, Eulophidae) including two new species. ZooKeys 154:9–30 doi:10.3897/zookeys.154.2158

Shevtsova E, Hansson C, Janzen DH, Kjærandsen J (2011) Stable structural color patterns displayed on transparent insect wings. Proceedings of the National Academy of Sciences of the United States of America 108:668–673 108:668–673 doi:10.1073/pnas.1017393108

Simon E (2013) Preliminary study of wing interference patterns (WIPs) in some species of soft scale (Hemiptera, Sternorrhyncha, Coccoidea, Coccidae). ZooKeys 319:269–281 doi:10.3897/zookeys.319.4219

Slatkin M (1984) Ecological Causes of Sexual Dimorphism. Evolution 38:622–630 doi:10.2307/2408711

Tang Y, Horikoshi M, Li W (2016) ggfortify: Unified interface to visualize statistical result of popular R packages. The R Journal 8 https://journal.r-project.org/.

Taylor LA, Cook C, McGraw KJ (2019) Variation in activity rates may explain sex-specific dorsal color patterns in Habronattus jumping spiders PLoS One 14:e0223015 doi:10.1371/journal.pone.0223015

Taylor PD, Merriam G (1995) Wing morphology of a forest damselfly is related to landscape structure. Oikos 73:43–48 doi:10.2307/3545723

Trischler C, Kern R, Egelhaaf M (2010) Chasing behavior and optomotor following in free-flying male blowflies: flight performance and interactions of the underlying control systems. Frontiers in Behavioral Neuroscience 4:20–20 doi:10.3389/fnbeh.2010.00020

Troje N (1993) Spectral categories in the learning behaviour of blowflies. Zeitschrift für Naturforschung C 48:96–104 doi:10.1515/znc-1993-1-218

Troscianko J, Stevens M (2015) Image calibration and analysis toolbox– a free software suite for objectively measuring reflectance, colour and pattern. Methods in Ecology and Evolution 6:1320–1331 doi:10.1111/2041-210X.12439

Uetz GW, Smith EI (1999) Asymmetry in a visual signaling character and sexual selection in a wolf spider. Behavioral Ecology and Sociobiology 45:87–93 doi:10.1007/s002650050542

Van Hateren J, Hardie R, Rudolph A, Laughlin S, Stavenga D (1989) The bright zone, a specialized dorsal eye region in the male blowfly Chrysomyia megacephala Journal of Comparative Physiology A 164:297–308 doi:10.1007/BF00612990

White TE, Vogel-Ghibely N, Butterworth NJ (2019) Flies exploit predictable perspectives and backgrounds to enhance iridescent signal salience and mating success. The American Naturalist doi:10.1086/707584

White TE, Zeil J, Kemp DJ (2015) Signal design and courtship presentation coincide for highly biased delivery of an iridescent butterfly mating signal. Evolution 69:14–25 doi:10.1111/evo.12551

Windig JJ, Nylin S (1999) Adaptive wing asymmetry in males of the speckled wood butterfly (*Pararge aegeria*)? Proceedings of the Royal Society B: Biological Sciences 266:1413–1418 doi:10.1098/rspb.1999.0795

Wootton RJ (1992) Functional morphology of insect wings. Annual Review of Entomology 37:113–140 doi:10.1146/annurev.en.37.010192.000553

