## Supplementary Material for "Love at first flight: Wing interference patterns are species-specific and sexually dimorphic in blowflies (Diptera: Calliphoridae)"

Table S1. Results of Permutation *F*-tests on RDA data from the RGB, CIELab and Blowfly datasets. Factors in bold are those that explained a significant proportion of overall WIP variation. Sp = species, Wi = wing.

| | | Sp | Sex | Wi | Sp:Sex | Sp:Wi | Sex:Wi | Sp:Sex:Wi |
|---|---|---|---|---|---|---|---|---|
| **RGB: colour** | *F* | 17.48 | 6.05 | 2.11 | 5.87 | 0.98 | 0.89 | 0.9 |
| | *P* | **0.001** | **0.001** | **0.047** | **0.001** | 0.459 | 0.529 | 0.584 |
| **RGB: contrast** | *F* | 44.24 | 7.28 | 2.11 | 3.57 | 1.2 | 1.18 | 0.78 |
| | *P* | **0.001** | **0.001** | 0.104 | **0.001** | 0.261 | 0.287 | 0.705 |
| **CIELab: colour** | *F* | 12.81 | 7.23 | 2.87 | 4.52 | 0.81 | 0.46 | 0.69 |
| | *P* | **0.001** | **0.001** | **0.01** | **0.001** | 0.817 | 0.919 | 0.933 |
| **CIELab: contrast** | *F* | 35.79 | 12 | 1.59 | 5.31 | 0.87 | 0.56 | 0.69 |
| | *P* | **0.001** | **0.001** | 0.156 | **0.001** | 0.605 | 0.662 | 0.817 |
| **Blowfly: colour** | *F* | 21.88 | 7.45 | 2.26 | 7.6 | 1.04 | 1.04 | 0.92 |
| | *P* | **0.001** | **0.001** | 0.064 | **0.001** | 0.414 | 0.37 | 0.515 |
| **Blowfly: contrast** | *F* | 25.22 | 7.24 | 2.25 | 6.49 | 1.26 | 1.45 | 0.79 |
| | *P* | **0.001** | **0.001** | 0.071 | **0.001** | 0.191 | 0.186 | 0.734 |

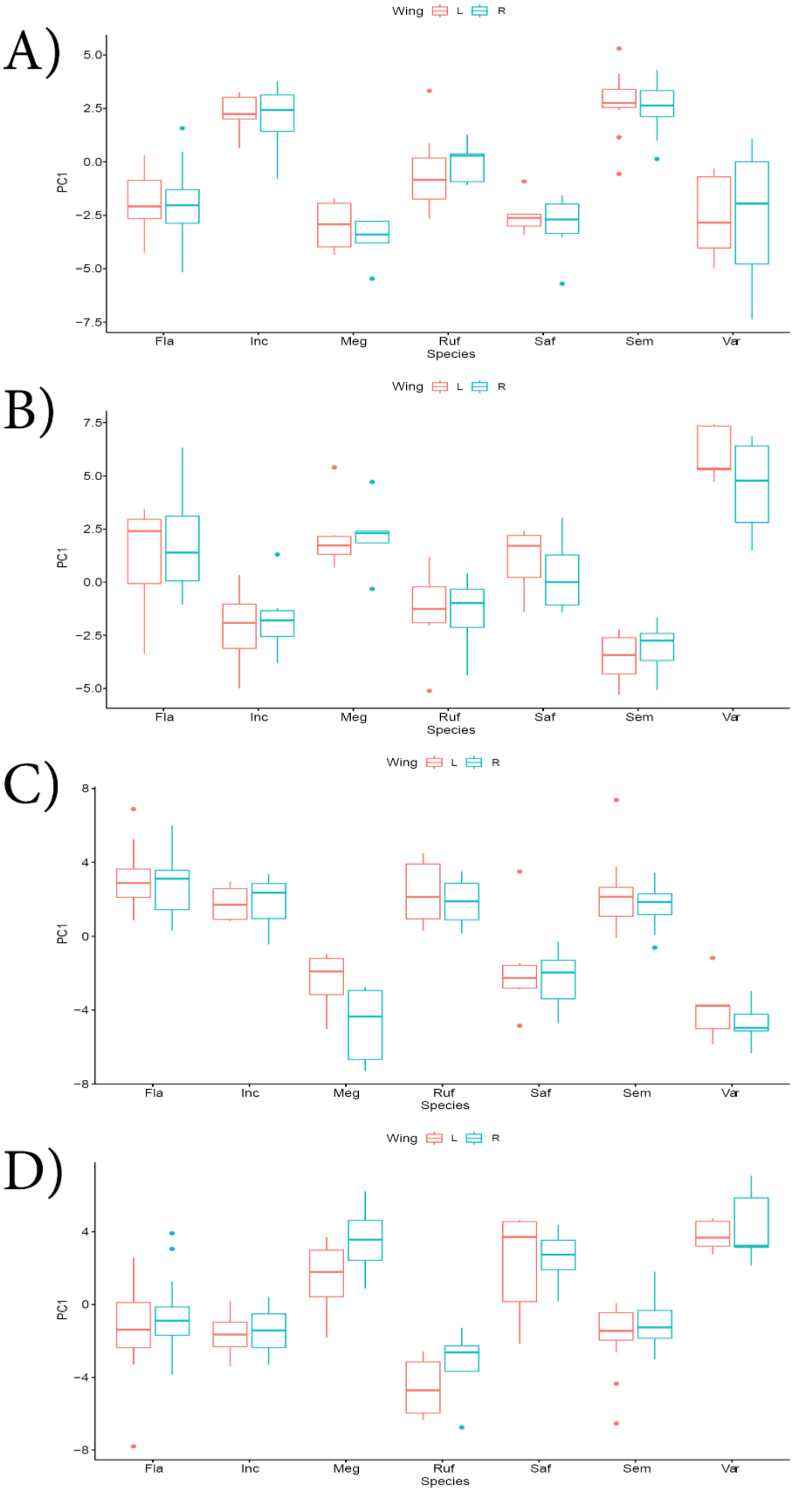


Figure S1. Asymmetry in WIPs represented by mean principal component (PC1) values for *Chrysomya* species (based on the blowfly dataset). A) female colour, B) female contrast, C) male colour, D) male contrast.

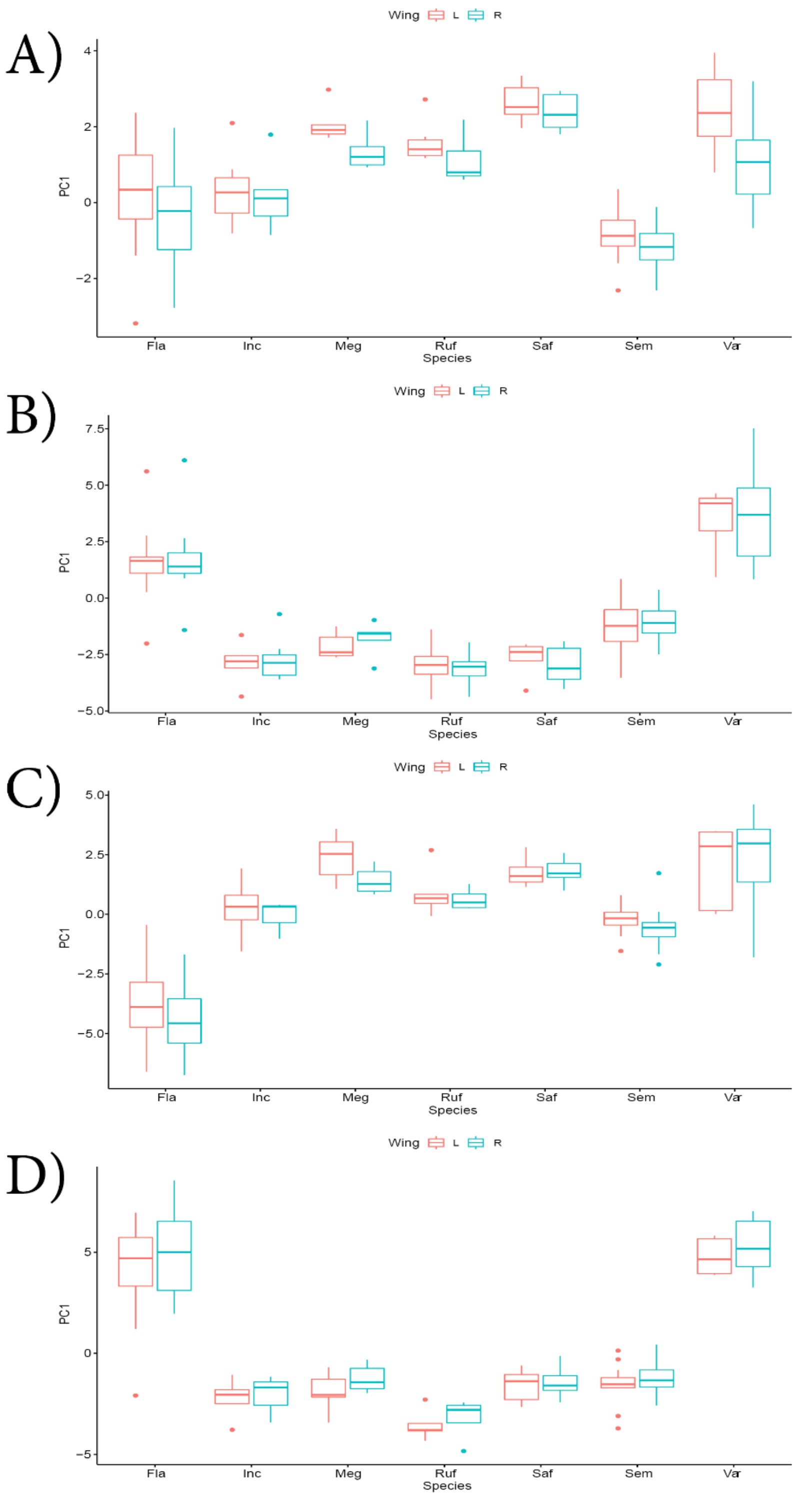


Figure S2. Asymmetry in WIPs represented by mean principal components (PC1) for *Chrysomya* species (based on the CIELab dataset). A) female colour, B) female contrast, C) male colour, D) male contrast.

**Figure S1 & S2 – Asymmetry in *Chrysomya* WIPs**

Importantly, while the RDA suggested that the effect of wing (left or right) was not significant when considered with species, sex, or species × sex (Supplementary Material 1: Table 1) inspection of mean PCA values (Supplementary Figures 1 & 2) suggests that there are intra-sexual differences in mean WIP colour and WIP colour contrast between left and right wings. Asymmetries in wing morphology have been widely reported in flying insects (Windig and Nylin 1999; Koshio et al. 2007; McLachlan 2010) and it is therefore likely that asymmetries in WIPs are also widespread. It is important that future studies consider asymmetries between left and right wings when assessing WIP variation – particularly considering that the symmetry of the WIP may itself be a component of the signal (Møller and Pomiankowski 1993; Uetz and Smith 1999).

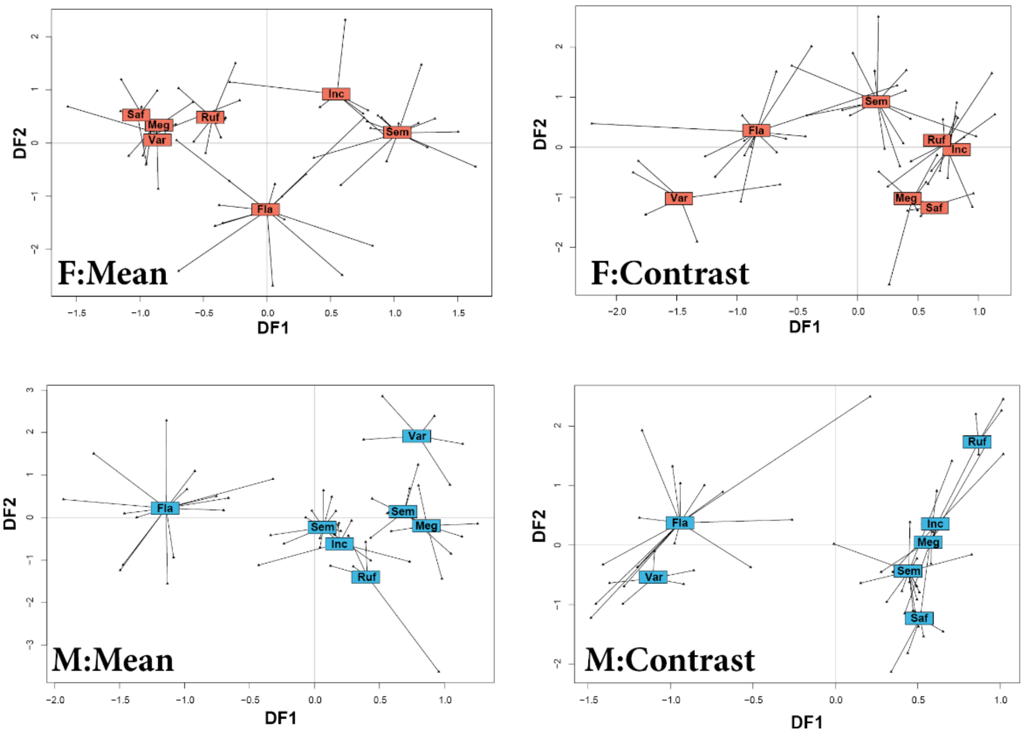


Fig S3. Quantitative differences in the wing interference patterns (WIP) of male (M) and female (F) Australian *Chrysomya* represented by discriminant factors 1 (DF1) and 2 (DF2). Results are from a redundancy discriminant analysis of WIP colour (as represented by average measurements of L, A and B values) and WIP colour contrast (as represented by standard deviations in L, A and B values). All measurements were made in CIELab 'human visual space' using the Multispectral Image Analysis and Calibration Toolbox for ImageJ (MICA toolbox) (Troscianko et al. 2019).

Table S2. Pairwise comparisons between species, based on redundancy discriminant analysis of WIP colour (as represented by average measurements of L, A and B values) and WIP colour contrast (as represented by standard deviations L, A and B values). All measurements were made in CIELab 'human visual space' using the Multispectral Image Analysis and Calibration Toolbox for ImageJ (MICA toolbox) (Troscianko et al. 2019). Bold values indicate significant differences. F = Female, M = Male.

| **F_Colour_CIELab** | Fla | Inc | Meg | Ruf | Saf | Sem |
|---|---|---|---|---|---|---|
| Inc | **0.0019** | - | - | - | - | - |
| Meg | **0.0019** | **0.0019** | - | - | - | - |
| Ruf | **0.0019** | **0.003** | 0.151 | - | - | - |
| Saf | **0.0019** | **0.003** | 0.588 | 0.0893 | - | - |
| Sem | **0.0019** | 0.0056 | **0.0019** | **0.0019** | **0.0019** | - |
| Var | **0.003** | **0.0019** | 0.982 | 0.2123 | 0.61 | **0.0019** |
| **F_Contrast_CIELab** | **Fla** | **Inc** | Meg | Ruf | Saf | **Sem** |
| Inc | **0.0023** | **-** | - | - | - | **-** |
| Meg | **0.0023** | **0.0292** | - | - | - | **-** |
| Ruf | **0.0023** | **0.951** | 0.0398 | - | - | **-** |
| Saf | **0.0023** | **0.0126** | 0.839 | 0.0161 | - | **-** |
| Sem | **0.0076** | **0.0126** | 0.0023 | 0.0161 | 0.0023 | **-** |
| Var | **0.0105** | **0.0023** | 0.0042 | 0.0023 | 0.0126 | **0.0023** |
| **M_Colour_CIELab** | **Fla** | **Inc** | Meg | Ruf | Saf | **Sem** |
| Inc | **0.0135** | **-** | - | - | - | **-** |
| Meg | **0.0035** | **0.1447** | - | - | - | **-** |
| Ruf | **0.0035** | **0.03832** | 0.0472 | - | - | **-** |
| Saf | **0.0035** | **0.1038** | 0.449 | 0.0336 | - | **-** |
| Sem | **0.0035** | **0.2354** | 0.009 | 0.0129 | 0.0129 | **-** |
| Var | **0.0035** | **0.0129** | 0.0105 | 0.0129 | 0.0129 | **0.0035** |
| **M_Contrast_CIELab** | **Fla** | **Inc** | Meg | Ruf | Saf | **Sem** |
| Inc | **0.0191** | **-** | - | - | - | **-** |
| Meg | **0.009** | **0.672** | - | - | - | **-** |
| Ruf | **0.007** | **0.0875** | 0.0372 | - | - | **-** |
| Saf | **0.0042** | **0.066** | 0.0766 | 0.0191 | - | **-** |
| Sem | **0.0042** | **0.0766** | 0.1659 | 0.0042 | 0.0766 | **-** |
| Var | **0.1116** | **0.0158** | 0.0163 | 0.0227 | 0.0042 | **0.0042** |

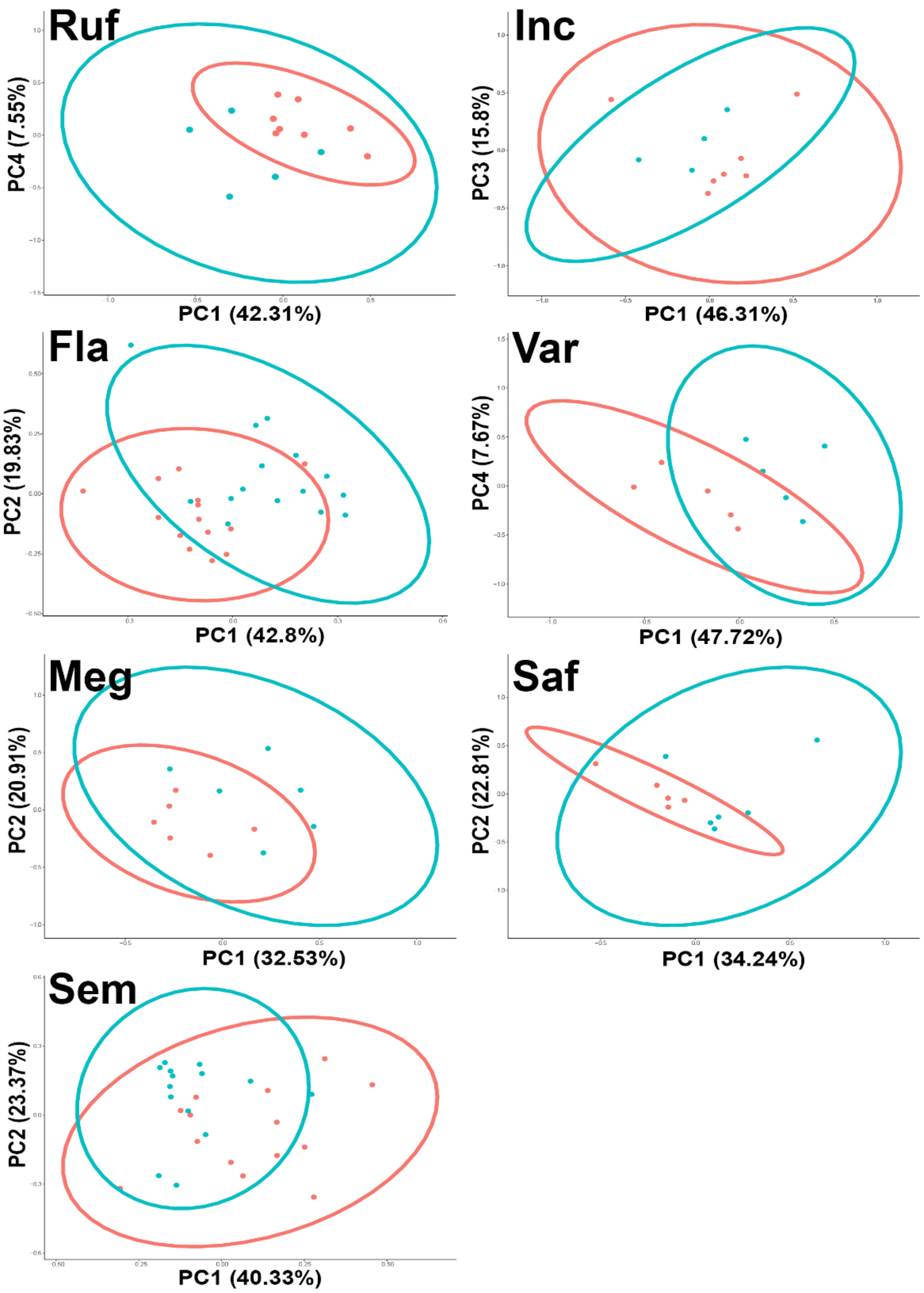


Figure S4. PCA plots of sex-specific differences in the CIELab average colour of WIPs (mean L, A and B values). The blue dots and ellipses represent males, while red dots and ellipses represent females.

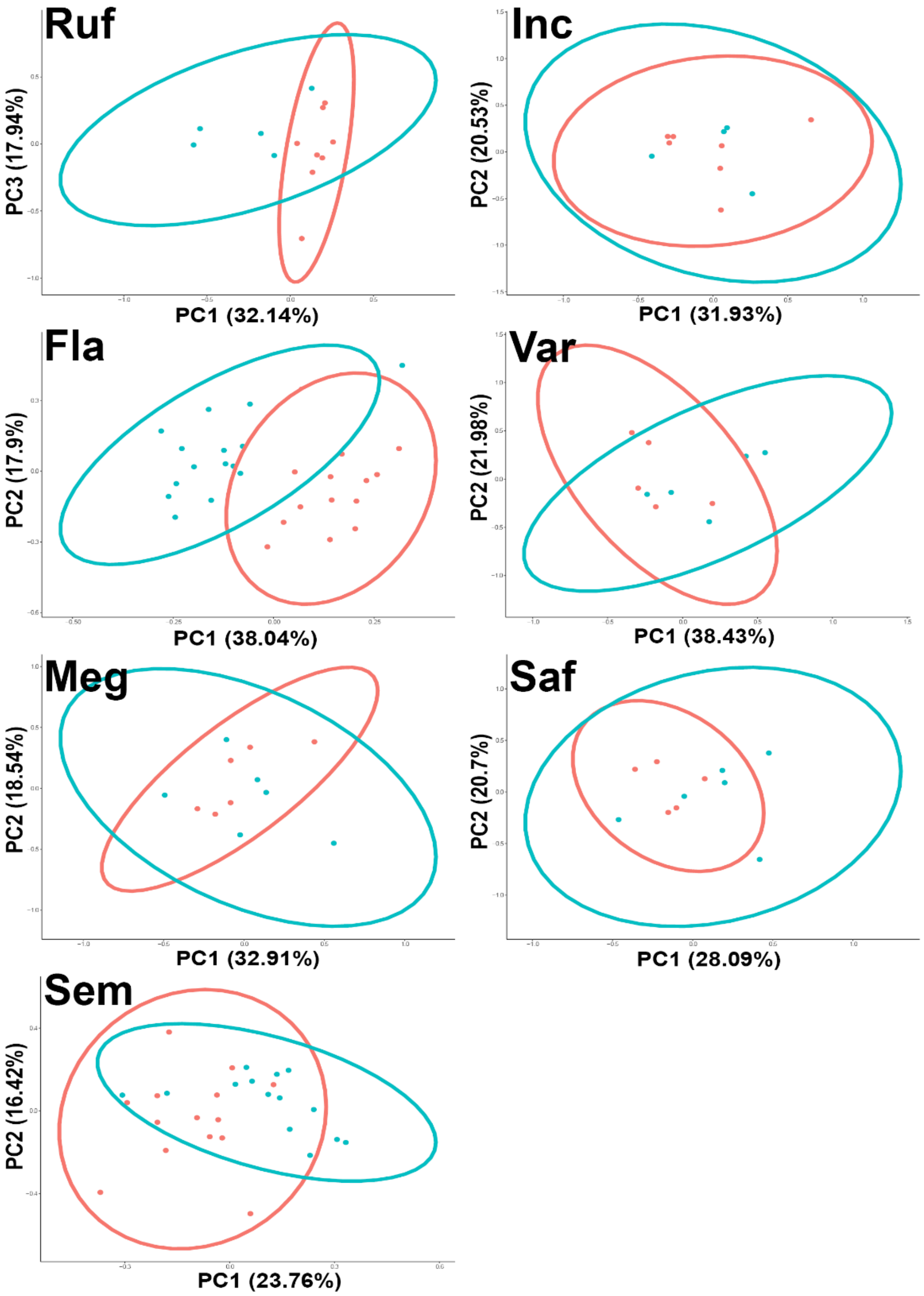


Figure S5. PCA plots of sex-specific differences in the CIELab average colour contrast of WIPs (mean L, A and B values). The blue dots and ellipses represent males, while red dots and ellipses represent females.

Table S3. a) Cumulative variances of principal components 1 – 5 from the blowfly colour dataset, that were used in ANOVA analysis to assess the differences in WIPs between the sexes of *Chrysoma* species. The output of ANOVA is also presented. Numbers in bold represent significant values.

| **Species (Calliphora)** | - | PC1 | PC2 | PC3 | PC4 | PC5 |
|---|---|---|---|---|---|---|
| Ruf | Cumulative variance (%) | 59.27 | 78.37 | 93.54 | 97.72 | 99.23 |
| | *F1,11* | 13.688 | 0.7813 | 0.0419 | 0.0167 | 3.6987 |
| | *P* | **0.003503** | 0.3957 | 0.8415 | 0.8996 | 0.08071 |
| Inc | Cumulative variance (%) | 46.13 | 71.84 | 84.44 | 95.69 | 98.77 |
| | *F1,9* | 0.49 | 1.4005 | 2.0142 | 0.5098 | 2.9931 |
| | *P* | 0.5016 | 0.2669 | 0.1895 | 0.4933 | 0.1177 |
| Fla | Cumulative variance (%) | 53.2 | 76.69 | 91.64 | 95.38 | 97.57 |
| | *F1,27* | 94.377 | 0.4477 | 0.3072 | 1.6691 | 0.3961 |
| | *P* | **2.63E-10** | 0.5091 | 0.584 | 0.2073 | 0.5344 |
| Var | Cumulative variance (%) | 56.96 | 77.02 | 88.39 | 94.06 | 97 |
| | *F1,8* | 1.5248 | 0.011 | 0.0599 | 2.4854 | 1.6594 |
| | *P* | 0.2519 | 0.9189 | 0.8128 | 0.1536 | 0.2337 |
| Meg | Cumulative variance (%) | 42.67 | 66.42 | 82.98 | 96.55 | 99.12 |
| | *F1,10* | 0.7832 | 0.2662 | 2.6287 | 0.438 | 5.9649 |
| | *P* | 0.3969 | 0.6171 | 0.136 | 0.5231 | **0.03471** |
| Saf | Cumulative variance (%) | 39.45 | 60.31 | 77.24 | 93.24 | 98.45 |
| | *F1,9* | 1.6813 | 2.4974 | 2.767 | 0.0154 | 0.0997 |
| | *P* | 0.227 | 0.1485 | 0.1306 | 0.9038 | 0.7594 |
| Sem | Cumulative variance (%) | 48.87 | 67.73 | 82.09 | 89.04 | 93.93 |
| | *F1,26* | 3.7301 | 3.1037 | 0.0804 | 1.1231 | 5.6487 |
| | *P* | 0.06441 | 0.08987 | 0.7791 | 0.299 | **0.02512** |

Table S3. b) The top five variables from the blowfly colour dataset that contributed significantly to the principal components. Contribution of variables was assessed using the function 'fviz_contrib' of the R package 'factoextra'.

| **Species** | **PC** | **Contributing variable** | **Wing Section** |
|---|---|---|---|
| *Ch. flavifrons* | 1 | Rh5 | E |
| | | Rh1 | E |
| | | Rh1 | Wing |
| | | Rh5 | Wing |
| | | Rh6 | E |
| *Ch. rufifacies* | 1 | Rh1 | B |
| | | Rh6 | B |
| | | Rh5 | B |
| | | Rh6 | Wing |
| | | Rh1 | Wing |
| *Ch. megacephala* | 5 | Rh6 | C |
| | | Rh5 | B |

| | | | |
|---|---|---|---|
| | | Rh5 | A |
| | | Rh1 | B |
| | | Rh5 | D |
| *Ch. semimetallica* | 5 | Rh6 | B |
| | | Rh1 | B |
| | | Rh5 | C |
| | | Rh1 | C |
| | | Rh5 | B |

Table S4. a) Cumulative variances of principal components 1 – 5 from the blowfly colour contrast dataset, that were used in ANOVA analysis to assess the differences in WIPs between the sexes of *Chrysoma* species. The output of ANOVA is also presented. Numbers in bold represent significant values.

| Species (Calliphora ) | - | PC1 | PC2 | PC3 | PC4 | PC5 |
|---|---|---|---|---|---|---|
| Ruf | Cumulative variance (%) | 61.34 | 80.79 | 93.22 | 97.78 | 99.17 |
| | *F1,11* | 14.202 | 1.9241 | 0.039 | 3.4238 | 0.0259 |
| | *P* | **0.003108** | 0.1929 | 0.847 | 0.09914 | 0.875 |
| Inc | Cumulative variance (%) | 54.03 | 76.79 | 88.52 | 95.09 | 97.47 |
| | *F1,9* | 0.2679 | 0.4512 | 0.3119 | 0.0018 | 2.3449 |
| | *P* | 0.6172 | 0.5186 | 0.5902 | 0.9667 | 0.1601 |
| Fla | Cumulative variance (%) | 49.16 | 69.78 | 81.97 | 89 | 92.62 |
| | *F1,27* | 20.712 | 25.799 | 0.0239 | 0.0527 | 0.0461 |
| | *P* | **0.0001017** | **2.46E-05** | 0.8782 | 0.8201 | 0.8315 |
| Var | Cumulative variance (%) | 41.55 | 71.01 | 82.79 | 90.17 | 95.49 |
| | *F1,8* | 16.985 | 0.3321 | 0.9219 | 0.2024 | 0.7569 |
| | *P* | **0.003339** | 0.5083 | 0.3651 | 0.6647 | 0.4088 |
| Meg | Cumulative variance (%) | 46.15 | 74.62 | 89.29 | 94.89 | 98.7 |
| | *F1,10* | 0.1532 | 1.3602 | 3.5836 | 0.4455 | 0.7338 |
| | *P* | 0.7037 | 0.2706 | 0.08762 | 0.5196 | 0.4117 |
| Saf | Cumulative variance (%) | 41.01 | 61.42 | 81.43 | 91.4 | 95.92 |
| | *F1,9* | 1.4182 | 0.9643 | 1.3615 | 0.5976 | 0.011 |
| | *P* | 0.2642 | 0.3518 | 0.2733 | 0.4593 | 0.9186 |
| Sem | Cumulative variance (%) | 53.8 | 67.31 | 80.01 | 87.23 | 91.2 |
| | *F1,26* | 17.254 | 2.9485 | 1.7786 | 0.0101 | 0.3729 |
| | *P* | **0.00003127** | 0.09785 | 0.1939 | 0.9208 | 0.5467 |

Table S4. b) The top five variables from the blowfly colour contrast dataset that contributed significantly to the principal components. Contribution of variables was assessed using the function ‘fviz_contrib’ of the R package ‘factoextra’.

| Species | PC | Contributing variable | Wing Section |
|---|---|---|---|
| *Ch. flavifrons* | 1 | Rh1 | C |
| | | Rh1 | B |
| | | Rh6 | C |

| | | | |
|---|---|---|---|
| | | Rh5 | B |
| | | Rh5 | C |
| | 2 | Rh6 | D |
| | | Rh6 | Wing |
| | | Rh5 | D |
| | | Rh6 | A |
| | | Rh5 | E |
| *Ch. rufifacies* | 1 | Rh1 | B |
| | | Rh6 | B |
| | | Rh6 | Wing |
| | | Rh1 | Wing |
| | | Rh5 | B |
| *Ch. varipes* | 1 | Rh1 | Wing |
| | | Rh6 | B |
| | | Rh1 | B |
| | | Rh6 | Wing |
| | | Rh1 | E |
| *Ch. semimetallica* | 1 | Rh5 | B |
| | | Rh1 | B |
| | | Rh6 | C |
| | | Rh1 | C |
| | | Rh1 | D |

Table S5. a) Cumulative variances of principal components 1 – 5 from the CIELab colour dataset, that were used in ANOVA analysis to assess the differences in WIPs between the sexes of *Chrysoma* species. The output of ANOVA is also presented. Numbers in bold represent significant values.

| **Species (CIELab)** | - | PC1 | PC2 | PC3 | PC4 | PC5 |
|---|---|---|---|---|---|---|
| Ruf | Cumulative variance (%) | 32.14 | 55 | 72.93 | 84.5 | 92.44 |
| | *F1,11* | 14.109 | 0.013 | 1.1298 | 0.8213 | 0.3927 |
| | *P* | **0.003176** | 0.9113 | 0.3106 | 0.3842 | 0.5437 |
| Inc | Cumulative variance (%) | 31.93 | 52.46 | 71.55 | 83.3 | 93.45 |
| | *F1,9* | 0 | 0.0109 | 2.6286 | 1.0388 | 4.7262 |
| | *P* | 0.9952 | 0.919 | 0.1394 | 0.3347 | 0.05774 |
| Fla | Cumulative variance (%) | 38.04 | 55.95 | 70.43 | 82.22 | 88.8 |
| | *F1,27* | 36.351 | 4.9144 | 0.4727 | 2.7435 | 1.2737 |
| | *P* | **1.96E-06** | **0.03524** | 0.4976 | 0.1092 | 0.269 |
| Var | Cumulative variance (%) | 38.43 | 60.41 | 80.62 | 88.16 | 93.49 |
| | *F1, 8* | 3.6538 | 0.1869 | 2.2759 | 2.2981 | 0.114 |
| | *P* | 0.09232 | 0.6769 | 0.1698 | 0.168 | 0.7443 |
| Meg | Cumulative variance (%) | 32.91 | 51.45 | 69.14 | 81.49 | 90 |
| | *F1,10* | 0.0892 | 0.7994 | 0.9076 | 0.0367 | 0.037 |
| | *P* | 0.7712 | 0.3923 | 0.3632 | 0.8519 | 0.8513 |
| Saf | Cumulative variance (%) | 28.09 | 48.79 | 65.73 | 82.07 | 91.35 |

| | | | | | | |
|---|---|---|---|---|---|---|
| | *F1,9* | 2.7562 | 0.3101 | 0.3086 | 4.736 | 1.3029 |
| | *P* | 0.1312 | 0.5912 | 0.5921 | 0.05753 | 0.2832 |
| Sem | Cumulative variance (%) | 23.76 | 40.18 | 53.96 | 67.12 | 76.89 |
| | *F1,26* | 12.535 | 0.2643 | 0.4679 | 0.4679 | 0.4383 |
| | *P* | **0.00153** | 0.2642 | 0.4679 | 0.4383 | 0.1629 |

Table S5. b) The top five variables from the CIELab colour dataset that contributed significantly to the principal components. Contribution of variables was assessed using the function 'fviz_contrib' of the R package 'factoextra'.

| Species | PC | Contributing variable | Wing Section |
|---|---|---|---|
| *Ch. flavifrons* | 1 | B | Wing |
| | | B | A |
| | | B | B |
| | | B | E |
| | | L | E |
| | 2 | A | Wing |
| | | B | D |
| | | L | B |
| | | B | E |
| | | A | B |
| *Ch. rufifacies* | 1 | L | B |
| | | B | B |
| | | L | Wing |
| | | B | E |
| | | B | C |
| *Ch. semimetallica* | 1 | B | C |
| | | B | E |
| | | B | D |
| | | B | B |
| | | A | E |

Table S6. a) Cumulative variances of principal components 1 – 5 from the CIELab colour contrast dataset, that were used in ANOVA analysis to assess the differences in WIPs between the sexes of *Chrysoma* species. The output of ANOVA is also presented. Numbers in bold represent significant values.

| Species (CIELab) | - | PC1 | PC2 | PC3 | PC4 | PC5 |
|---|---|---|---|---|---|---|
| Ruf | Cumulative variance (%) | 42.31 | 65.09 | 79.39 | 86.94 | 91.4 |
| | *F1,11* | 5.1646 | 0.0309 | 0.7712 | 3.8139 | 0.3058 |
| | *P* | **0.04411** | 0.8636 | 0.3986 | 0.07674 | 0.5913 |
| Inc | Cumulative variance (%) | 46.31 | 66.83 | 82.63 | 88.76 | 93.64 |
| | *F1,9* | 0.8551 | 1.4801 | 0.1619 | 3.9265 | 0.0192 |
| | *P* | 0.3792 | 0.2546 | 0.6968 | 0.07886 | 0.8928 |
| Fla | Cumulative variance (%) | 42.8 | 62.63 | 74.35 | 81.84 | 87.01 |

| | | | | | | |
|---|---|---|---|---|---|---|
| | *F1,27* | 13.304 | 7.2756 | 0.0259 | 5.2265 | 4.8251 |
| | *P* | **0.001116** | **0.0119** | 0.8734 | **0.03031** | **0.03861** |
| Var | Cumulative variance (%) | 47.72 | 65.67 | 80.39 | 88.07 | 92.56 |
| | *F1,8* | 13.459 | 0.8668 | 0.2908 | 0.2954 | 0.9589 |
| | *P* | **0.006322** | 0.8668 | 0.2908 | 0.2954 | 0.9589 |
| Meg | Cumulative variance (%) | 32.53 | 53.44 | 67 | 79.61 | 88.08 |
| | *F1,10* | 6.3463 | 2.2419 | 0.3124 | 0.259 | 0.3228 |
| | *P* | **0.03043** | 0.1652 | 0.5885 | 0.6218 | 0.5825 |
| Saf | Cumulative variance (%) | 34.24 | 57.05 | 68.81 | 79.63 | 87.26 |
| | *F1,9* | 7.7448 | 0.0857 | 0.3196 | 0.2323 | 1.9014 |
| | *P* | **0.0213** | 0.7763 | 0.5856 | 0.6413 | 0.2012 |
| Sem | Cumulative variance (%) | 40.33 | 63.7 | 73.3 | 80.29 | 86.65 |
| | *F1,26* | 7.1634 | 4.6796 | 0.7039 | 0.2001 | 0.6233 |
| | *P* | **0.01271** | **0.0399** | 0.4091 | 0.6584 | 0.4369 |

Table S6. b) The top five variables from the CIELab colour contrast dataset that contributed significantly to the principal components. Contribution of variables was assessed using the function 'fviz_contrib' of the R package 'factoextra'.

| Species | PC | Contributing variable | Wing Section |
|---|---|---|---|
| *Ch. flavifrons* | 1 | B | Wing |
| | | B | A |
| | | B | D |
| | | B | B |
| | | B | C |
| | 2 | L | E |
| | | L | B |
| | | L | C |
| | | A | C |
| | | A | A |
| *Ch. varipes* | 1 | B | D |
| | | A | C |
| | | B | Wing |
| | | B | A |
| | | L | B |
| *Ch. rufifacies* | 1 | A | Wing |
| | | A | B |
| | | L | C |
| | | B | Wing |
| | | B | C |
| *Ch. megacephala* | 1 | A | Wing |
| | | A | A |
| | | B | D |
| | | A | C |
| | | A | D |

| | | | |
|---|---|---|---|
| *Ch. saffranea* | 1 | A<br>A<br>A<br>B<br>L | B<br>A<br>D<br>D<br>B |
| *Ch. semimetallica* | 1 | B<br>A<br>B<br>A<br>A | A<br>B<br>D<br>C<br>D |
| | 2 | L<br>B<br>L<br>L<br>B | B<br>E<br>D<br>C<br>C |
